# *LRRK2* G2019S variant is associated with transcriptional changes in Parkinson’s disease human myeloid cells under proinflammatory environment

**DOI:** 10.1101/2024.05.27.594821

**Authors:** Elisa Navarro, Anastasia G. Efthymiou, Madison Parks, Giulietta M Riboldi, Ricardo A. Vialle, Evan Udine, Benjamin Z. Muller, Jack Humphrey, Amanda Allan, Charlie Charalambos Argyrou, Katia de Paiva Lopes, Alexandra Münch, Deborah Raymond, Rivka Sachdev, Vicki L. Shanker, Joan Miravite, Viktoryia Katsnelson, Katherine Leaver, Steve Frucht, Susan B Bressman, Edoardo Marcora, Rachel Saunders-Pullman, Alison Goate, Towfique Raj

## Abstract

The G2019S mutation in the leucine-rich repeat kinase 2 (LRRK2) gene is a major risk factor for the development of Parkinson’s disease (PD). LRRK2, although ubiquitously expressed, is highly abundant in cells of the innate immune system. Given the importance of central and peripheral immune cells in the development of PD, we sought to investigate the consequences of the G2019S mutation on microglial and monocyte transcriptome and function. We have generated large-scale transcriptomic profiles of isogenic human induced microglial cells (iMGLs) and patient derived monocytes carrying the G2019S mutation under baseline culture conditions and following exposure to the proinflammatory factors IFNγ and LPS. We demonstrate that the G2019S mutation exerts a profound impact on the transcriptomic profile of these myeloid cells, and describe corresponding functional differences in iMGLs. The G2019S mutation led to an upregulation in lipid metabolism and phagolysosomal pathway genes in untreated and LPS/IFNγ stimulated iMGLs, which was accompanied by an increased phagocytic capacity of myelin debris. We also identified dysregulation of cell cycle genes, with a downregulation of the E2F4 regulon. Transcriptomic characterization of human-derived monocytes carrying the G2019S mutation confirmed alteration in lipid metabolism associated genes. Altogether, these findings reveal the influence of G2019S on the dysregulation of the myeloid cell transcriptome under proinflammatory conditions.

## Introduction

Parkinson’s disease (PD) is a progressive neurodegenerative disorder with no cure and no approved disease-modifying treatments. PD is highly heritable, and early studies of PD focused on causal genes implicated in familial PD including *SNCA, GBA, and LRRK2*^1–4^. Recently, genome-wide association studies (GWAS) have identified 90 independent genetic loci across 78 genomic regions that contribute to sporadic forms of the disease, and approximately 15% of people with PD, but without risk variants in PD associated genes, have family history of PD in first-degree relatives. These findings provide compelling evidence for the existence of a heritable component in sporadic PD^5^. We have previously demonstrated that PD risk alleles are expression quantitative trait loci (eQTLs) in peripheral monocytes and that monocytes and microglia derived from PD patients show profound transcriptional dysregulation^6–8^. Data from other groups have also shown an enrichment of PD heritability in immune cell types, suggesting that PD risk genes play a major role within these cell types^6–12^.

GWAS have identified PD-associated loci harboring genes known to be involved in familial PD, indicating a potential shared molecular basis between sporadic and familial forms of the disease.^13^ *LRRK2* is one such gene, with common and rare variants that confer a modest to high risk for developing PD^8,13–15^. These *LRRK2* variants have been shown to be associated with a host of alterations in many cellular processes including disordered vesicular trafficking, cytoskeletal dynamics, mitochondrial function, autophagy and lysosomal function, as well as immune system dysregulation^16^. While numerous single nucleotide polymorphisms (SNPs) have been discovered within the *LRRK2* locus, the most prevalent variant associated with PD involves a Glycine to Serine substitution at amino acid position 2019 (G2019S), which is within the kinase domain and is thought to increase kinase activity^17^. *LRRK2* G2019S-linked PD (LRRK2-PD) exhibits clinical features that are indistinguishable from idiopathic PD, has increasing penetrance with age, and accounts for 1-5% of sporadic cases of PD and 1-18% of familial cases depending on ancestral population^18^. Due to the genetic overlap between LRRK2-PD and other familial PD, as well as the similar clinical presentation of patients between LRRK2-PD and idiopathic PD, studying the role of LRRK2 in PD is critical both for understanding disease etiology as well as developing targets for therapeutic intervention.

LRRK2 is ubiquitously expressed within the brain, and much research has focused on the role of the G2019S mutation on neuronal and astrocytic function^16^. However, the highest LRRK2 expression is seen in immune cells, particularly peripheral blood mononuclear cells (PBMCs), and this gene seems to play a pivotal role in immune regulation. LRRK2 has been shown to be associated with the activation and maturation of human monocytes^19^ and in modulating neuroinflammation by cytokine signaling^20,21^. Variants in *LRRK2* have been implicated in different immune-related disorders including Crohn’s disease and leprosy^22^, or cancer^23^, while elimination of the *LRRK2* gene seems to suppress immune system activation^21^. *LRRK2* risk variants have been reported to be associated with higher levels of LRRK2 expression in monocytes^8^ and monocyte-derived microglia^24^. Moreover, *LRRK2* expression is highly dependent on inflammatory state^19,24–27^, and has been shown to respond to different proinflammatory stimuli such as interferon-γ (IFNγ)^25^ ^27^, lipopolysaccharides (LPS)^28^, IFNβ, tumor necrosis factor α (TNFα), and interleukin-6 (IL-6)^19^. Finally, it has also been reported that non-coding common genetic variants influence *LRRK2* expression specifically in microglia, pointing to the cellular specificity of genetic variation and highlighting the role of microglia in the context of *LRRK2*^29^.

Collectively, these data suggest that there may be an association between the innate immune system and PD, especially in PD patients carrying mutations in *LRRK2*. Due to the reduced penetrance of disease onset in LRRK2 carriers, we hypothesize that *LRRK2* variants affect innate immune cells, such as monocytes, macrophages, and microglia, in proinflammatory conditions to affect disease onset and progression. Thus, our objective is to characterize and understand the transcriptional and functional effect of the G2019S variant in myeloid cells. Here, we have used human induced microglial cells as well as patient-derived monocytes and characterized the G2019S effect in untreated cells and following stimulation with the proinflammatory regulators IFNγ and LPS. Our findings demonstrate that the G2019S mutation has a strong influence in myeloid cells transcriptome and functionality, affecting phagocytosis, lipid metabolism, and proliferation genes.

## Results

### Generation of isogenic *LRRK2* G2019S iPSC-derived microglia

We investigated the impact of proinflammatory stimuli in microglia, brain-resident macrophages that express LRRK2 and are implicated in PD. To reduce variability and enhance statistical power with small sample size, we utilized an isogenic induced pluripotent stem cell (iPSC) line differentiated to induced microglial cells (iMGLs). We obtained three isogenic CRISPR-edited iPSC clonal lines from a single donor, one carrying wild-type (WT), one carrying heterozygous (het), and one carrying homozygous (hom) *LRRK2* G2019S through the iPSC Neurodegenerative Disease Initiative (iNDI)^30^. We differentiated these iPSCs to iMGLs using a previously described protocol^31^ and treated them with IFNγ or LPS for 24 hours (**Fig. 1A**). Differentiated iMGLs express the microglial marker IBA1, as well as LRRK2 (**Fig. 1B**) and stimulation of WT iMGLs with IFNγ or LPS showed increased expression of *IFIT1*, *IL1B*, and *TNFa* compared to untreated WT iMGLs (**Fig. S1B**). IFNγ treatment also induced an increase in expression of LRRK2, which has also been previously described in the literature^19^. For downstream analyses, we collected data from three independent differentiations of each clonal line followed by 24 hours of treatment (untreated, LPS and IFNγ), yielding a total of 27 samples (**Supplementary Table 1**).

**Figure 1.**
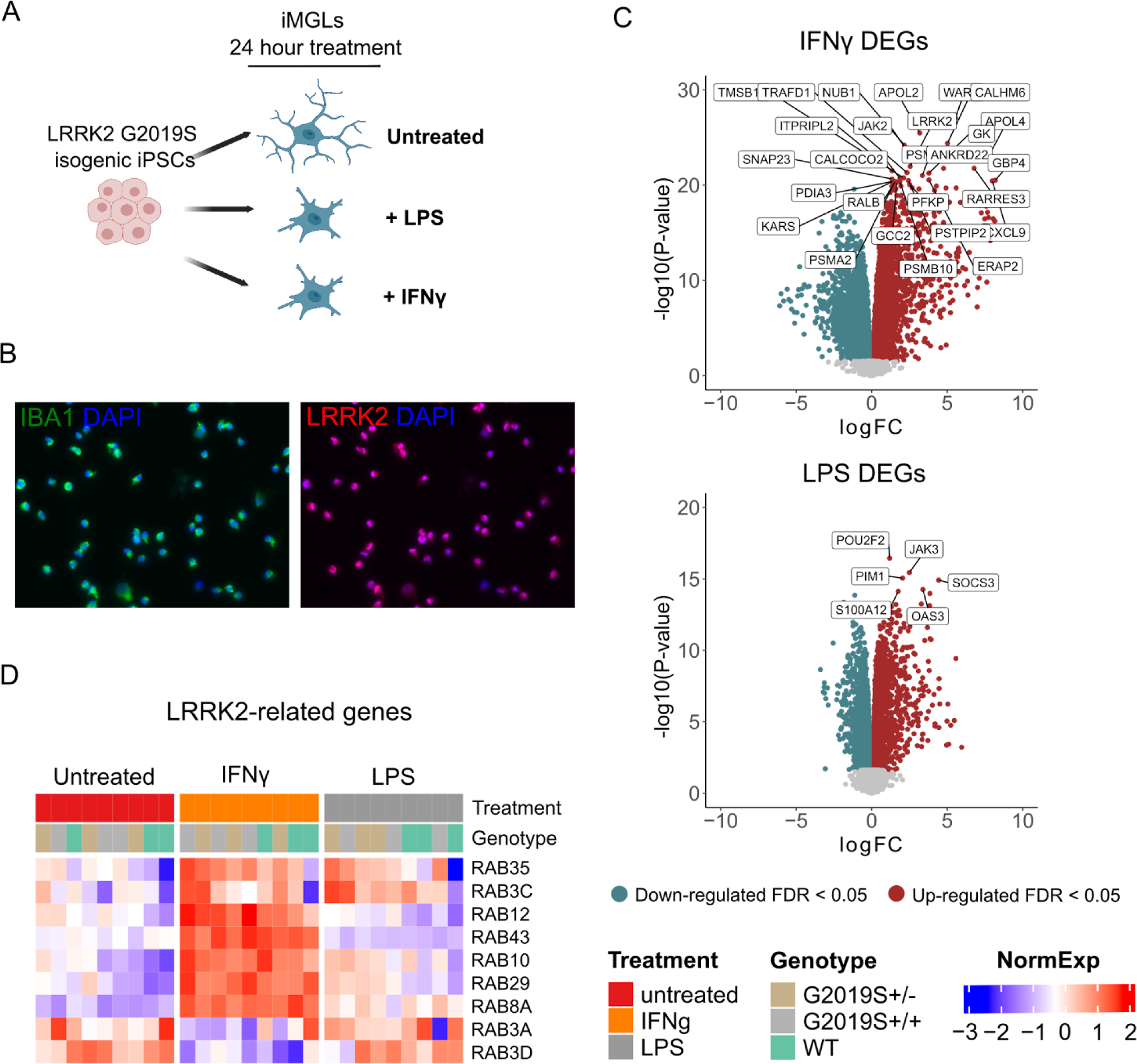
Human induced microglial cells respond to IFNγ and LPS. (A) Overview of the study design. CRISPR-edited isogenic induced pluripotent stem cells for the G2019S mutation in the *LRRK2* gene were differentiated to microglial cells (iMGLs) and stimulated with IFNγ or LPS for 24 hrs. (B) Representative immunofluorescence pictures of iMGLs showing the expression of the microglial marker IBA-1 and LRRK2 after differentiation. (C) Volcano plots showing differential expression after stimulation with IFNγ (top) or LPS (bottom). Plots show the FC of genes (x axis, log_2_ scale) and their *P* value significance (y axis, −log_10_ scale). Differentially expressed genes (DEGs) at FDR < 0.05 are highlighted in red (upregulated genes) and blue (downregulated genes). A moderated *t*-statistic (two-sided) was used as the statistical test. (D) Heatmap showing the normalized expression of a subset of *LRRK2*-interactors ^34^(y axis) across samples (x axis), clustered by treatment. Blue represents normalized expression < 0 and red represents normalized expression > 0.

### iMGLs respond to IFNγ and LPS with profound transcriptomic changes and cytokine release

We first investigated the effect of proinflammatory stimuli (IFNγ and LPS) on the iMGL transcriptome. To do that, we studied differences between samples treated with LPS or IFNγ in comparison to untreated samples, without considering genotype. After extensive quality control measures, we observed both proinflammatory stimuli induced a large change in the number of differentially expressed genes (DEGs) when compared to untreated samples (5955 for LPS and 7856 for IFNγ at FDR < 0.05; **Supplementary tables 2 and 3**), with a significant association of DEGs between both conditions (p-value = 2.2e-138, Odds ratio = 2.6, overlap tested using Fisher’s exact test)^32^(**Fig. 1C**). Moreover, we also assessed if changes in gene expression also reflected a change in the cytokine profiles of iMGLs. Stimulation with IFNγ and LPS induced a significant increase in the secretion of multiple proinflammatory cytokines as indicated in **Fig. S1D** and **Supplementary Table 4** (two-way ANOVA with Dunnett’s post hoc test, *P* < 0.05), including IL-12p40, IL-15, IL-1RA, and TNFa, which were increased by both conditions. These results demonstrate that iMGLs both transcriptionally and functionally respond to proinflammatory stimuli.

We then performed gene set enrichment analysis (GSEA) on predefined lists from Gene Ontology: Biological Processes (GO:BP) to identify enriched pathways in these DEGs and visualized the results using Cytoscape (**Fig. S1C**). The predominant auto-annotated clusters involved an upregulation of immune response pathways, as expected. Both treatments also induced a downregulation of protein localization, transcription, and translation pathways. We performed an enrichment analysis using a list of 407 genes that have been reported as potential interactors of LRRK2^33^. We found that both stimuli, IFNγ and LPS, induced a response which was significantly enriched for LRRK2 interactors (Fisher’s exact test, IFNγ *P* = 0.0146; LPS *P* = 0.00012). These results suggest that both stimuli induce transcriptional changes in genes closely associated with LRRK2 and therefore may indeed have disease relevant effects.

### Transcriptional dysregulation in G2019S iMGLs under proinflammatory stimuli

We performed RNA-sequencing on WT, G2019S^+/−^ and G2019S^+/+^ iMGLs, with and without IFNγ or LPS stimulation. In line with the notorious effect of G2019S on human health outcomes, we did observe a strong transcriptomic dysregulation when comparing heterozygous G2019S^+/−^ (het) or homozygous G2019S^+/+^ (hom) genotypes vs WT in untreated (717 DEGs in G2019S^+/−^ and 2692 DEGs in G2019S^+/+^) and stimulated iMGLs with LPS (587 DEGs in G2019S^+/−^ and 4009 DEGs in G2019S^+/+^) or IFNγ (410 DEGs in G2019S^+/−^ and 1389 DEGs in G2019S^+/+^). (**Fig. 2A, 2B, Supplementary Tables 5-10**). Among these genes, 91% of het DEGs were shared between het and hom comparisons in untreated iMGLs, 82% under IFNγ and 94% in LPS (**Fig. 2B**), indicating a dosage-dependent effect of the G2019S mutation on gene expression changes. In order to see if the G2019S mutation has a context-specific effect on myeloid cells, we studied the interaction between genotype and treatment. We obtained that 5 genes showed a significant interaction between IFNγ and G2019S mutation (*ITPRIP*, *AKAP2*, *CYSLTR2*, *CXorf21* & *SPOPL*) and 2 genes in LPS condition (*SEPT4* & A*C068946.2*) (**Supplementary Tables 11 & 12**).

**Figure 2.**
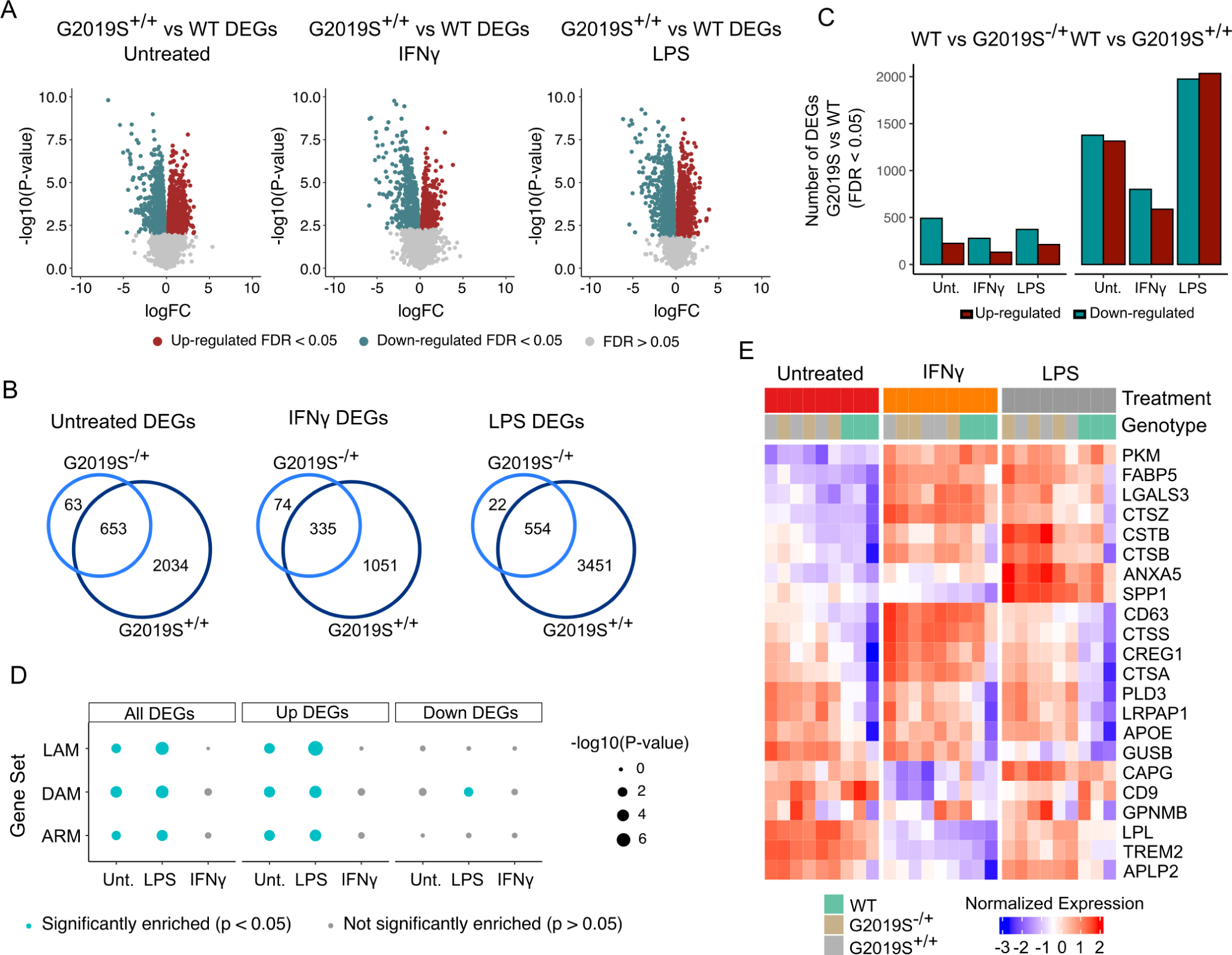
The G2019S mutation drives transcriptomic dysregulation in iMGLs upon IFNγ and LPS stimulation. (A) Volcano plots showing differential expression between G2019S^+/+^ and WT in untreated condition and after stimulation with IFNγ or LPS. Plots show the FC of genes (x axis, log_2_ scale) and their *P* value significance (y axis, −log_10_ scale). DEGs at FDR < 0.05 are highlighted in red (upregulated genes) and blue (downregulated genes). A moderated *t*-statistic (two-sided) was used as the statistical test. (B) Venn diagrams showing the number of shared DEGs between heterozygous (G2019S^+/−^) iMGLs and homozygous iMGLs (G2019S^+/+^) vs WT. (C) Barplot showing the number of DEGs between G2019S and WT iMGLs under different stimulations. Red bars represent the number of upregulated DEGs (FC > 0; FDR < 0.05) and blue bars represent the number of downregulated DEGs (FC < 0; FDR < 0.05). (D) Enrichment for microglial signatures in G2019S^+/+^iMGLs in untreated condition and upon stimulation. Blue dots represent microglial phenotypes for which there is a significant enrichment (*P* < 0.05; Fisher’s-exact test) in G2019S^+/+^transcriptome. DAM = damage-associated microglia; LAM = lipid-accumulating macrophages, ARM = activated response microglia. (E) Heatmap showing the normalized expression of a selection of DEGs in WT, G2019S^+/−^ and G2019S^+/+^. The heatmap shows a selection of DAM/LAM/ARM genes (y axis) across samples (x axis), clustered by genotype. Blue represents normalized expression < 0 and red represents normalized expression > 0.

We investigated whether G2019S^+/+^ iMGLs compared to WT exhibit a similar transcriptional profile to any known microglial/macrophages subpopulations described in the literature, such as disease/damage-associated microglia (DAM)^35–37^, activated response microglia (ARM)^38^, or lipid-accumulating macrophages (LAM)^39^. We performed enrichment analysis using gene lists for these microglia/macrophage phenotypes that can be found in **Supplementary Table 13**. We found that DEGs in G2019S^+/+^ untreated iMGLs and upon LPS stimulation were significantly enriched for a list of 110 DAM genes (Fisher’s exact test, Untreated: *P* = 1.16e-4; LPS: *P* = 8.03e-6) (**Fig. 2D**). We also found that upregulated DEGs driven by G2019S^+/+^ mutation were enriched for ARM genes^38^(list containing 131 genes), in untreated and LPS condition (Fisher’s exact test, Untreated: *P* = 0.01; LPS: *P* = 4.5e-4) (**Fig. 2D**). Whether this ARM response is specific to the amyloid plaque deposition model or general protein aggregation and/or other disease aspects remains to be determined. Finally, we also explored if the phenotype driven by G2019S^+/+^ could be reflecting alterations in lipid homeostasis. To do that, we used a LAM signature gene list and observed that the transcriptomic dysregulation in G2019S^+/+^ untreated and LPS stimulated iMGLs was also enriched for this macrophage signature (Fisher’s exact test, Untreated: *P* = 4.6e-3; LPS: *P* = 3.03e-6). This enrichment for different disease-associated phenotypes was more prominent in up-regulated DEGs, reflecting that G2019S mutation increases the expression of these signatures. These data indicate that *LRRK2* G2019S iMGLs exhibit a gene expression signature similar to that found in microglia/macrophages within other disease models.

### Stimulated G2019S^+/+^ iMGLs display an upregulation of phagosome and lipid metabolism genes

We performed Gene Set Enrichment Analysis (GSEA) using Gene Ontology (GO) terms on the DEGs between G2019S^+/+^ and WT iMGLs in the untreated condition and after stimulation with IFNγ and LPS to determine which biological pathways were most affected by this mutation. We observed a great number of GO:Biological Process terms among upregulated DEGs (**Supplementary Table 14**) with many DEGs enriched in phagocytosis and lipid metabolism pathways (**Fig. 3A**). Together, these data suggest that G2019S mutation could drive an increased expression in phagocytosis and lipid metabolism genes.

**Figure 3.**
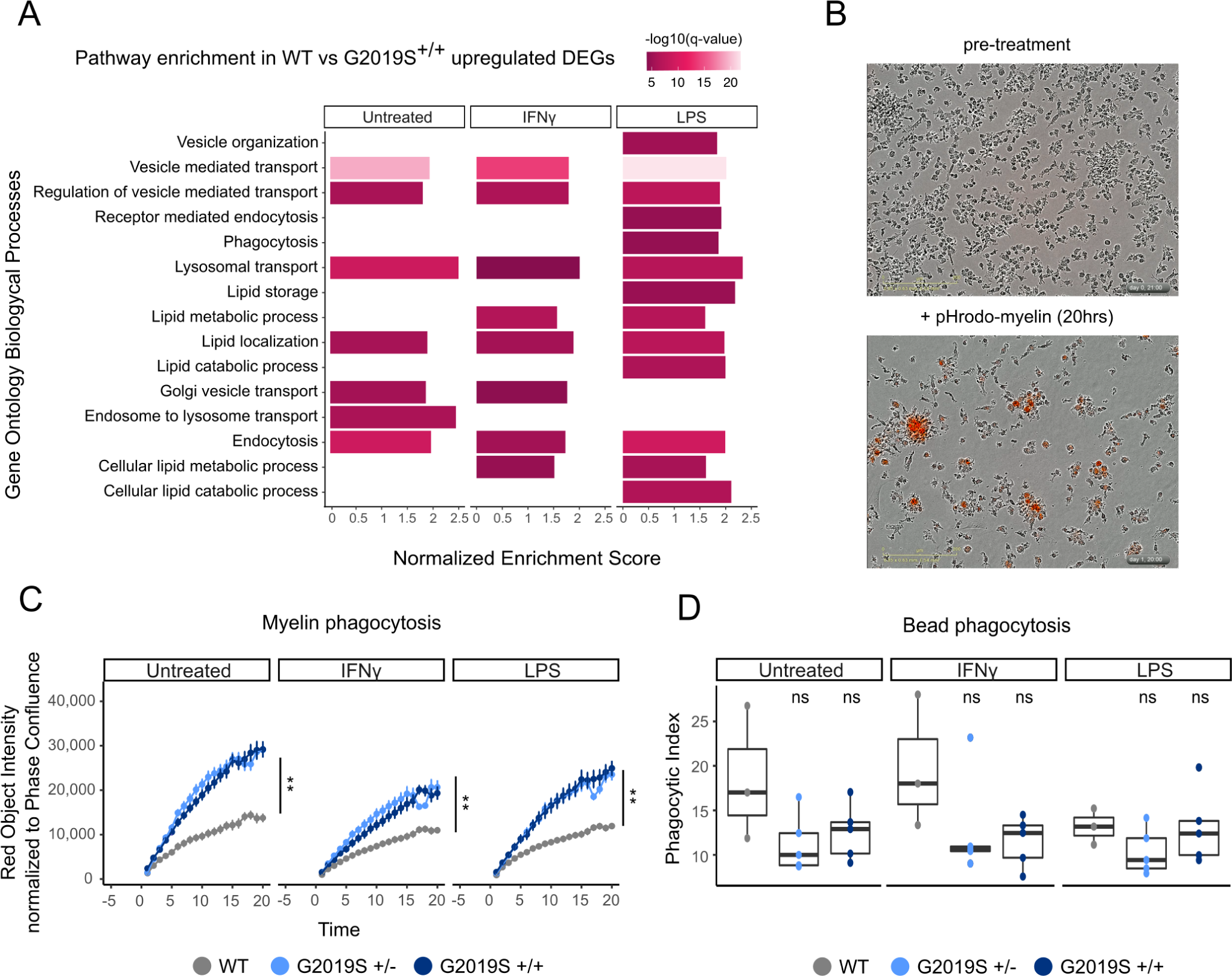
G2019S^+/+^ iMGLs show alterations in lipid metabolism and phago-lysosomal pathways. (A) Pathway enrichment analysis for the upregulated DEGs using Biological Process from GSEA. Pathways are represented in the y axis and normalized score in the x axis. Significance (-log_10_ *P* value scale of the q-value) is represented by color. Only significant pathways (q-value < 0.05) of pathways related to lipid metabolism and phagocytosis are represented. (B) Representative bright focus microphotographs of iMGLs subjected to ph-rodo labeled human myelin. Top picture shows iMGLs before treatment and bottom picture shows iMGLs after 20 hrs treatment with pH-rodo labeled human myelin. (C) Myelin phagocytosis of iMGLs under the different treatments measured by IncuCyte microscope. The y axis shows the phagocytic capacity measured as red object intensity normalized by cell confluence and the x axis the different time points. Genotypes begin to diverge in their myelin phagocytosis after 3 hours (n = 3 independent differentiations, Linear mixed effects model ** *P* value < 0.001) and show significant differences across genotypes by the end of 20 hours (n = 3 independent differentiations, ANOVA with Dunnett’s post-hoc test ** *P* value < 0.001). (D) Carboxylated-beads phagocytosis of iMGLs under the different treatments measured by FACS. Y axis represents the phagocytic index and x axis the different treatments (n = 3 independent differentiations, ANOVA with Dunnett’s post-hoc test).

To determine whether the transcriptional changes we observed have any functional implications, we performed a phagocytosis assay using human myelin debris, a lipid-rich substrate. iMGLs were treated with 25 µg of pHrodo-conjugated human myelin debris for 20 hours and imaged every hour to monitor the pHrodo fluorescent signal over time. G2019S^+/−^ and G2019S^+/+^ cells phagocytosed more myelin than their WT counterparts under both untreated and proinflammatory conditions (**Fig. 3B, C**). This finding remained consistent when measuring both the intensity of the red pHrodo signal and the total red object area. To determine whether this observed difference in phagocytosis is due to increased uptake of phagocytic substrates or a failure to clear those substrates, we treated iMGLs with carboxylated beads for 24 hours. Because these beads are autofluorescent, we could not use live imaging; instead, we measured their accumulation using flow cytometry. No significant differences in bead uptake were observed across genotypes or treatments, indicating that the observed dissimilarity in myelin phagocytosis between genotypes is unlikely to be attributed to general differences in substrate uptake (**Fig. 3D**). However, further investigation is required to comprehensively understand the effect of the G2019S mutation on the processes of uptake and degradation of lipid-rich debris such as myelin fragments. Together with the transcriptomic data, our results suggest that the difference in myelin phagocytic clearance is regulated by downstream processes within the endolysosomal system rather than phagocytic uptake.

### Stimulated G2019S iMGLs display a downregulation of mitosis and cell-cycle genes and decreased proliferation

We then explored mechanisms associated with downregulated DEGs in G2019S*^+/+^* compared to WT iMGLs, and we found a significant enrichment for cell cycle and mitosis pathways under proinflammatory stimuli (**Fig. 4A, Supplementary Table 14**). To determine whether G2019S^+/+^ iMGLs exhibited deficits in proliferation compared to WT iMGLs, we performed live imaging analysis using IncuCyte and measured phase object confluence of iMGLs over a 24 hour period (**Fig. 4B**). We observed that G2019S iMGLs showed a decreased rate of object confluence over that period, irrespective of treatment, suggesting decreased proliferation. While these data serve as an indirect measure of proliferation, it does support the notion that G2019S iMGLs display diminished proliferation compared to their wild-type counterparts.

**Figure 4.**
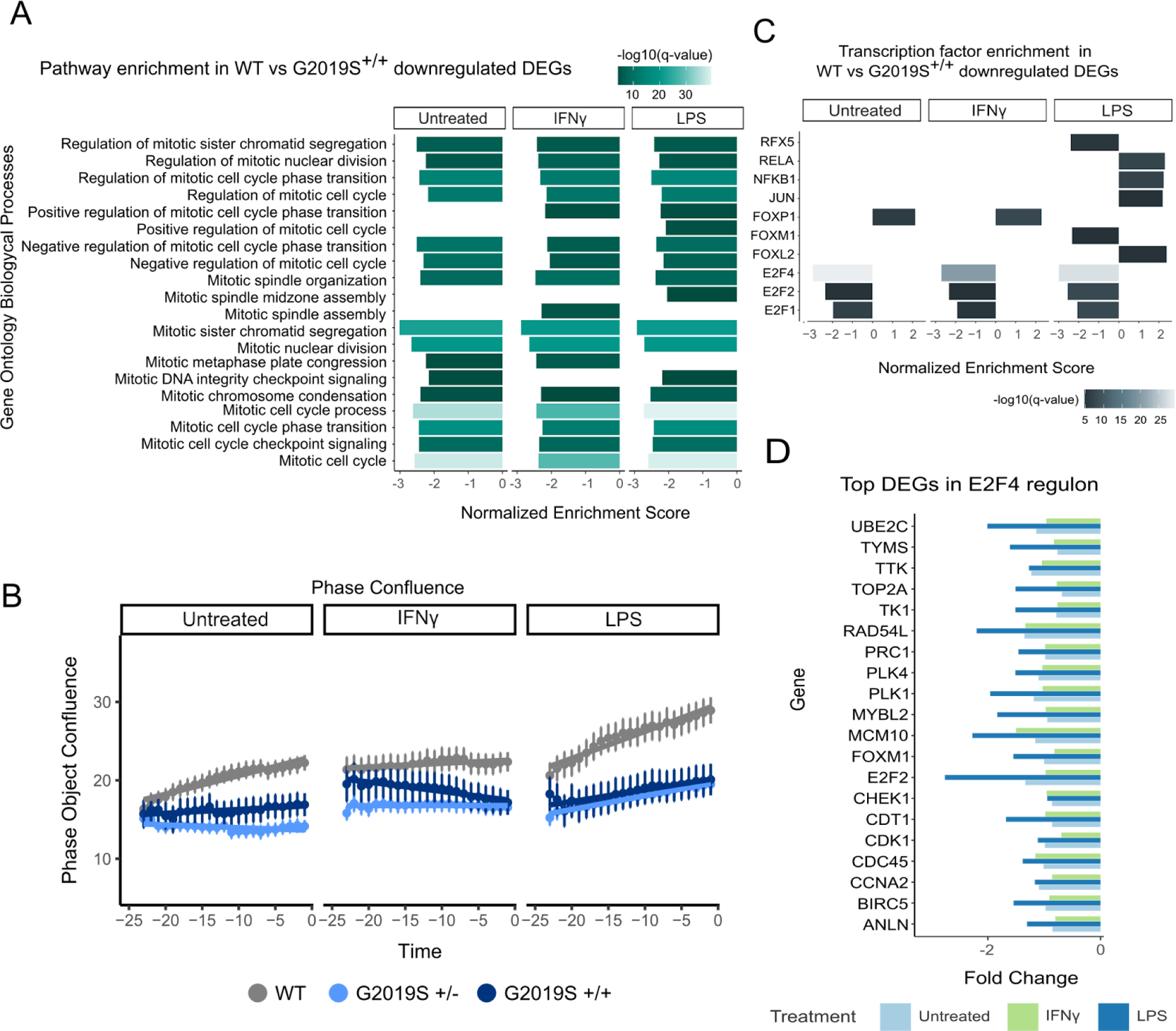
G2019S^+/+^iMGLs show downregulation in E2F regulon genes. (A) Pathway enrichment analysis for the downregulated DEGs using Biological Process from GSEA. Pathways are represented on the y axis and normalized score on the x axis. Significance (-log_10_ *P* value scale of the q-value) is represented by color. Only significant pathways (q-value < 0.05) related to cell mitosis and proliferation are represented. (B) Phase object confluence measurements of iMGLs using IncuCyte live-imaging. The y axis shows the phase object confluence and the x axis represents time, with samples separated by treatment. (C) Transcription factor enrichment using Dorothea. TFs are represented on the y axis and normalized enrichment score on the x axis. Significance (-log10 P value scale of the q-value) is represented by color. Only significant regulons (q-value < 0.01) are represented. (D) Boxplot showing fold change of the top 20 DEGs within the *E2F4* regulon under different treatments. Y axis shows gene names and x axis the fold change.

We used the regulatory target gene sets list from DoRothEA to identify whether there was an enrichment in particular transcription factor motifs in the G2019S^+/+^ iMGL DEGs that may be mediating the cell cycle and mitosis effects. Interestingly, we observed that G2019S^+/+^ displayed a strong downregulation in E2F transcription factor targets in both LPS and IFNγ proinflammatory treatment conditions (**Fig. 4C, Supplementary Table 15**). The E2F family of transcription factors is known to regulate cell proliferation and consists of both transcriptional activators and repressors^40^. The most significant enrichment was observed for E2F4 regulon in which we observed significant downregulation of its target genes without differential expression of E2F4 itself (**Fig. 4D**).

### G2019S mutation is associated with transcriptomic dysregulation in human derived monocytes

Thus far, our data suggest a profound effect of *LRRK2* G2019S genotype and proinflammatory stimuli on the iMGL transcriptome. In order to understand whether these *in vitro* findings from one isogenic iPSC pair are representative of changes observed in people with Parkinson’s we performed the same transcriptomic analysis in human monocytes. Monocytes were obtained from a total of 35 participants divided in 4 groups based on PD diagnosis and *LRRK2* mutation: PD cases without the G2019S mutation (PD-WT), PD cases with the G2019S mutation (PD-G2019S), healthy subjects without the *LRRK2* G2019S mutation (CTRL-WT), and non-PD manifesting carriers of the G2019S variant (CTRL-G2019S) All patients were age-matched and sex balanced (**Fig. S3A & B and Supplementary Table 16**). Following the same paradigm as before, we isolated monocytes, stimulated the cells with the proinflammatory stimuli LPS and IFNγ for 24h, and performed RNA-seq on this set of samples.

We first sought to confirm whether IFNγ or LPS stimulation induces similar transcriptional changes in human monocytes as in cultured iMGLs. We performed DE analysis irrespective of disease or *LRRK2* mutation genotype and observed that both stimuli, IFNγ and LPS, trigger a profound transcriptional response in human monocytes (**Fig. S5A, B, Supplementary Table 17 & 18**). Again, we observed that *LRRK2* expression was significantly increased after IFNγ, but did not change upon LPS stimulation (**Fig. S3C, D**), similar to what we observed in iMGLs and as has been described in the literature^27,41^. We did not observe any changes in the level of *LRRK2* expression in carriers vs non carriers either with or without PD (**Fig. S3C**). We then performed a similar enrichment analysis using curated lists of PD and LRRK2-interactor genes (**Supplementary Table 13**) and observed that both proinflammatory stimuli triggered profound changes in the expression of those genes (**Fig. S5C, D**). Overall, these results highlight that IFNγ and LPS can produce a proinflammatory response on the human monocyte transcriptome which is relevant in the context of PD.

We then investigated what effect the G2019S mutation had on gene expression in human monocytes. We performed DE between G2019S carriers and non carriers separately for each stimuli (baseline, IFNγ, and LPS). Although the number of significant DEGs was limited due to high variability and sample size (**Supplementary Tables 19-21**), we found an increase in the number of DEGs between carriers vs non carriers upon stimulation (**Fig. S6A, B**). This effect is also observed in the qq-plot (**Fig. 5A**). These results highlight that similar to what we observed in the *in vitro* iMGL experiments, the *LRRK2* G2019S effect on gene expression in human monocytes is stronger under proinflammatory conditions, linking together the genetic and environmental contribution to pathogenesis.

**Figure 5.**
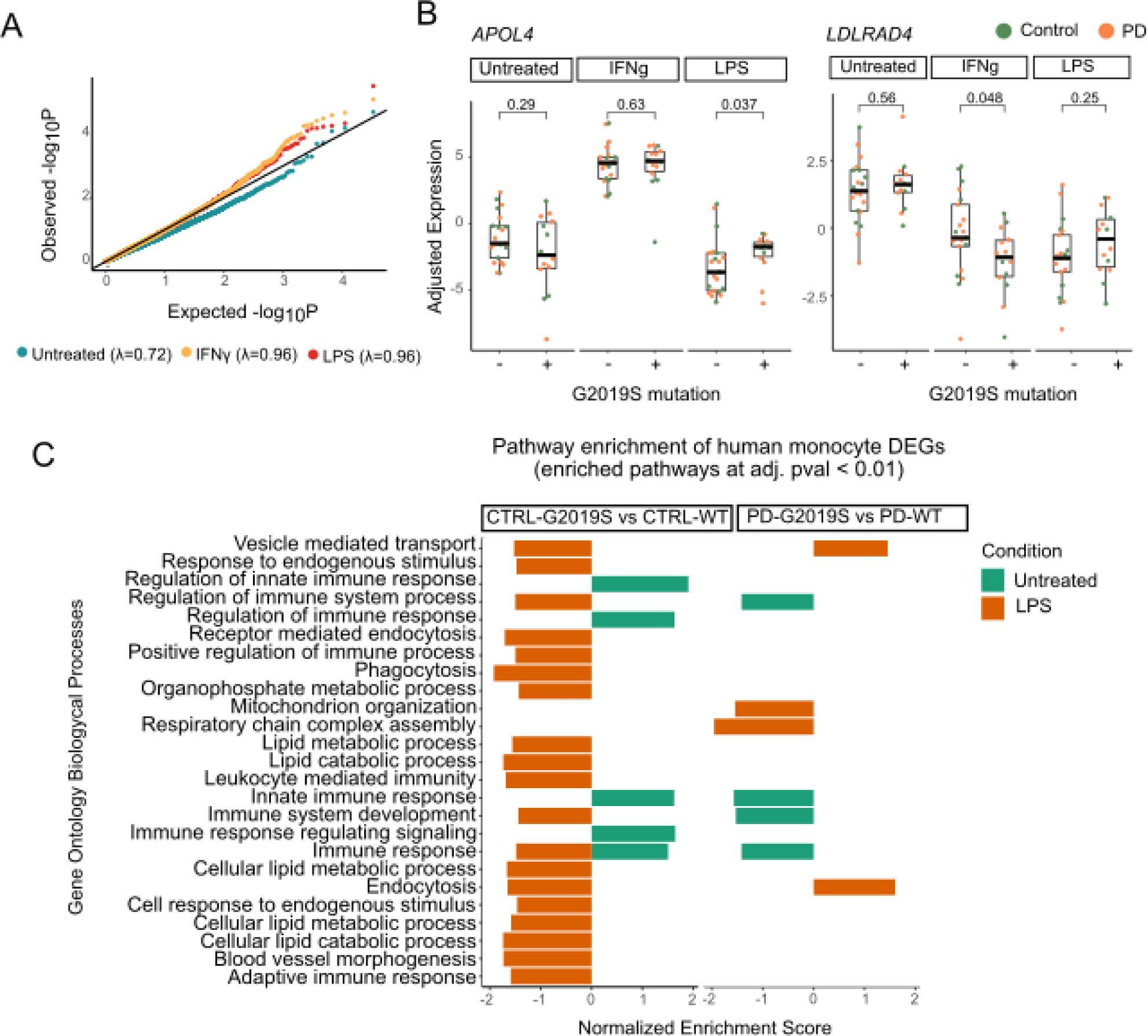
Patient derived monocytes carrying the G2019S mutation show transcriptomic dysregulation. (A) Quantile quantile (qq)-plot shows the observed distribution of *P* values which deviates from the expected uniform distribution across the three different treatments. The genomic inflation factor (λ or *X*^2^ median) was calculated as the measure of deviations of the observed distribution from the expected null distribution. (B) Examples of selected DEGs (*APOL4* on the left and *LDLRAD4* on the right) in which there is significant interaction between treatment and genotype. Adjusted expression of *voom* normalized counts after regression of covariates is shown. In boxplots, the line represents the median and boxes extend from the 25th to the 75th percentile. P-values calculated with Wilcoxon test. (C) Pathway enrichment analysis of G2019S carriers vs non carriers distributed by diagnosis, control on the left and PD on the right. Pathway enrichment was performed with a pre ranked approach from GSEA using Biological Processes. Y axis shows the significant pathways and x axis normalized expression score. Only significant pathways with q-value < 0.01 are shown.

To deepen the link of genetics and environment, we further explored the interaction between the G2019S mutation carrier status and the two proinflammatory treatments, and noticed statistically significant changes in several genes related to lipid metabolism, in line with the previous results shown in iMGLs (**Supplementary Tables 22 & 23**). An example is *APOL4*, a member of the apolipoprotein L family which plays a role in lipid exchange. We demonstrated that *APOL4* expression is significantly upregulated in G2019S carriers only under LPS but not under the other conditions (**Fig. 5B**), which links the proinflammatory environment and the mutation status as both co-regulators of the expression of this gene. Another example is the *LDLRAD4* gene, a member of the low-density lipoprotein receptors which has been involved in TGF-beta signaling. This gene is significantly downregulated in G2019S carriers under IFNγ stimulation, but not in baseline or LPS (**Fig. 5B**). These examples illustrate context specific gene regulation in human monocytes, demonstrating that the G2019S mutation affects gene expression in a context specific manner.

### Human monocytes carrying the G2019S mutation also display transcriptomic alterations in lipid metabolism and phagocytosis related genes

We further sought to understand the effect of G2019S in the context of health and disease. G2019S is one major risk factor to develop PD, so understanding the transcriptional signature among carriers with PD vs healthy individuals could give insights into the pathogenesis. To approach this question we performed DE analysis comparing the effect of G2019S mutation within each of the diagnoses performing the following analyses: (i) CTRL-G2019S vs CTRL-WT and (ii) PD-G2019S carriers vs PD-WT. A major observation is that proinflammatory stimuli induce a higher number of DEGs between the different diagnoses (PD or Controls) rather than in untreated condition (**Supplementary Tables 24-29**). In order to better understand the transcriptomic dysregulation associated with G2019S mutation in the different groups, we performed pathway analyses using GO:BP, as described previously (**Fig. 5C, Supplementary Table 30**). Surprisingly, even though the number of significant DEGs was significantly increased under stimulation, in the case of IFNγ we did not observe significant enrichment for any specific biological process. In untreated samples we observed enrichment in a number of pathways related to innate immune response. Nevertheless, it is noteworthy that under LPS stimulation we observed a significant enrichment in pathways related to phagocytosis and endocytosis, as well as lipid metabolic processes. These results are of importance as they reinforce the observations made in iMGLs, showing how they capture the biological effects of G2019S mutation, which are translatable to human primary myeloid cells. They also reinforce the effect of *LRRK2* in regulating phagocytosis and lipid metabolism in a context specific manner. This transcriptional dysregulation is orchestrated by a number of TFs highlighted in **Fig. S6C**, including *STAT1* and *STAT2* (implicated in the immune response) or *SPI1* and *FOXP1* (implicated in the myeloid differentiation), whose downstream genes are enriched upon G2019S mutation carriers compared to controls (**Supplementary Table 31**). All together, we demonstrate how *LRRK2* G2019S affects the monocyte transcriptome, especially under proinflammatory stimulation, reinforcing the role of this gene in myeloid cell function and the innate immune response.

Finally, we also explored the transcriptomic alterations driving disease within the LRRK2-G2019S carriers, we compared dysregulation between PD-G2019S vs CTRL-G2019S. Results from differential expression analysis in untreated monocytes and under stimulation are included in **Supplementary Tables 32-34**. Pathway analysis revealed that monocytes from symptomatic patients present down-regulation in pathways associated with immune response and defense. They also show a down-regulation in genes related to lipid metabolism and endocytosis and vesicle trafficking (**Fig S7, Supplementary Table 35**). Nevertheless, when monocytes are stimulated with either IFNγ or LPS, there is swift in directionality, with pathways related to immune response, lipid metabolism, or vesicle transport being upregulated in PD-G2019S vs CTRL-G2019S after proinflammatory stimuli treatment. Although further experiments are needed, these results show a differential response to proinflammatory stimuli of PD vs CTRL monocytes among donors with the G2019S mutation, demonstrating that LRRK2 may be mediating the response of monocytes to these proinflammatory stimuli.

## Discussion

*LRRK2* G2019S is a common PD mutation that has been shown to impact phagocytosis, autophagy, inflammation, vesicular trafficking, mitophagy, and lysosomal function in various neural cells^15^. Despite the high expression of *LRRK2* in immune cells, its role in myeloid cells has been underexplored. Given the implication of microglia and other innate immune cells in the pathogenesis of PD, we sought to investigate the effect of *LRRK2* G2019S and proinflammatory signals on the human microglial and monocyte transcriptome. For this study, we combined iPSC-derived microglial cells as well as human isolated monocytes carrying the G2019S mutation in order to shed light into the functional effect of *LRRK2* gene in myeloid cells. Recently, Ohtonen and collaborators conducted a similar approach, generating iPSC-derived microglia from PD donors with the G2019S mutation^42^. Our findings corroborate several of their observations, and incorporate additional treatments and analyses to further expand on the interaction between genotype and environmental stimuli. The robustness of both studies is further enhanced by our incorporation of human-derived monocytes from PD patients and age-matched controls, both with and without the G2019S mutation. This broader, population-based approach reinforces the observations made using isogenic iPSCs, offering a more comprehensive view of the disease mechanism. Through our iMGL modeling, we observed that the transcriptomic alterations seen in our G2019S^+/+^ clones were enriched for several disease-associated microglial signatures, including DAM, LAM, and ARM. These results show that G2019S iMGLs are capturing the transcriptomic network of different disease associated microglial phenotypes related to neurodegenerative diseases^43^.

Large scale genetic studies link genes associated with lipid metabolism with the risk of PD, including *GBA*, *ASAH1*, *SREBF1*, *GALC* and *DGKQ*. These findings corroborate our observations, as G2019S microglia and patient derived monocytes under LPS stimulation display dysregulation in pathways related to lipid metabolism. Indeed, human plasma and cerebrospinal fluid of *LRRK2*-G2019S carriers has been reported to present important alterations in lipid metabolites^44^ and postmortem brain tissues show alpha-synuclein colocalized with lipid-rich structures^45^. These human patient-based observations are in line with increasing evidence in animal models showing the importance of lipids in regulating microglial functionality and how its dysregulation may contribute to neurodegeneration^46,47^. In parallel, we also observed a profound dysregulation in phago-lysosomal genes in G2019S^+/+^ cells. Several studies have also explored the role of G2019S on phagocytosis, showing that this mutation increases myeloid cell phagocytosis of various substrates including beads and *E. coli* particles^48,49^ and *LRRK2* inhibition has been shown to decrease microglia engulfment of beads and neuronal axons^50^. The previous studies speculate that *LRRK2* G2019S’s increased phosphorylation of downstream targets including the WAVE complex and RAB proteins, which regulate phagosome and endosome fusion, may be mediating the increase in myeloid cell phagocytosis. The alteration in phagolysosomal genes, combined with changes in lipid transport, storage, and metabolism pathways, were reflected in the functional data that we obtained, showing increased accumulation of lipid-rich myelin debris in G2019S iMGLs. Additionally, our results noted that the increase in myelin accumulation by G2019S iMGLs was not accompanied by an increased bead uptake, which is in line with the observation of Ohtonen and collaborators using iPSC *LRRK2*-G2019S derived microglia^42^, and also similar to other studies^51^, suggesting that the increased myelin accumulation may be due to a defect in clearing the lipid debris. It is possible that the increased phosphorylation activity of *LRRK2*-G2019S is also having an immediate effect on phagocytosis-related proteins in iMGLs, nevertheless more experiments that focus on these downstream phosphorylation targets could help to decipher the molecular mechanism. This could be in line with recent findings demonstrating that monocyte derived microglia from PD patients with the G2019S mutation also showed altered capability to transfer aggregates of alpha synuclein, and are more vulnerable to cytotoxicity^52^. Moreover, a recent paper identified 25 protein QTLs (pQTLs) in cerebrospinal fluid associated with variants in *LRRK2*^53^ which are related to phagolysosomal trafficking. Indeed, 6 of the 25 proteins identified in that paper are also differentially expressed in our G2019S iMGLs under LPS stimulation (*CD68, FCGR1A, FTL, GAA, GRN, TMEM106A*), all 6 being upregulated in our G2019S^+/+^ clones.

We also found that downregulated DEGs in G2019S^+/+^iMGLs showed association with mitosis, cell cycle, and proliferation pathways. Many of the downregulated genes were enriched for E2F transcription factor binding motifs, suggesting a mechanism by which this downregulation may be occurring. E2Fs are a family of transcription factors that are involved in cell cycle regulation, and consist of both transcriptional activators and repressors. The downregulation of these E2F target genes suggests that the repressive E2Fs are being influenced by LRRK2. E2F4 and its binding partners can be phosphorylated, and this phosphorylation regulates E2F4 localization and activity^54^. Previous studies have postulated increased expression of E2Fs as a mechanism responsible for neuronal loss in PD^55,56^, while E2F1 ablation could confer neuroprotection^57^. Nevertheless, the function of this family of transcription factors seems to be cell specific, as in astrocytes increased E2Fs is postulated to favor proliferation in response to injury^58^, as a mechanism of plasticity. The implication of this pathway in myeloid cells has been less explored. Our study links *LRRK2*-G2019S and cell cycle disruption in human microglia. The role of this proliferative phenotype remains unclear. A recent paper by Belhocine et al. uncovered two context-specific programs of transcriptional regulation of microglial proliferation *in vivo*, the latter of which involved E2F transcription factors as a regulatory checkpoint to enable microglia to enter the cell cycle^59^. Although it is clear that more research is needed to determine the function of these different E2F family members in microglia and whether these cell cycle disruptions contribute to the pathogenicity of *LRRK2*-G2019S in PD, we could hypothesize that the downregulation of these factors along with the lower proliferative capacity could lead to lack of adaptation to brain injury in PD.

Our study was designed to deepen our understanding of the transcriptional and functional alterations in monocytes and microglia carrying the G2019S mutation, though it did contain several limitations. This study was conducted using isogenic cells from a single donor and it is therefore important to note that our observations could reflect clonal differences. We attempted to address this limitation by integrating human derived monocytes from carriers and non carriers of the *LRRK2*-G2019S mutation, as a means of adding population heterogeneity to our study. The inclusion of this population-based approach lends support to our conclusions, as it yields similar themes in the results. Our current study also does not take into account haplotype contributions; *LRRK2*-G2019S has been found on three major haplotypes, and polygenic risk score has been shown to affect *LRRK2* PD penetrance in affected individuals^60^. However, these genetic interactions are beyond the current scope of our study. Notwithstanding our study’s focus on a single donor line, the inclusion of both heterozygous and homozygous *LRRK2*-G2019S iMGLs, along with three independent differentiation batches, allowed us to investigate whether there was a dose-dependent effect of genotype on the microglial transcriptome. And finally, while this study mainly focused on transcriptional dysregulation, future studies with increased sample size will also focus on functional approaches (including lipidomics and proliferation assays) to decipher how the pathways identified may be contributing to Parkinson’s pathology.

In summary, this study revealed that G2019S mutation in the *LRRK2* gene produces transcriptomic dysregulation in human microglia and monocytes under proinflammatory environments, and that the main pathways altered are related to phago-endosomal processes, lipid metabolism, and cell division. These results demonstrate the link between genetics and environment, and highlight the importance of myeloid cells in PD pathogenesis. Future studies will be important to deepen our understanding of how *LRRK2* regulates these cellular pathways in the immune system and to which extent contributes to the onset and progression of Parkinson’s disease.

## Methods

### iPSC culture

Isogenic iPSCs for G2019S (wild-type, heterozygous, and homozygous clones from one donor) were provided by the Cookson Lab (NIH)^30^. Genotype was validated by Sanger sequencing, and all clones displayed a normal karyotype and standard pluripotent expression markers. iPSCs were maintained in StemFlex medium and cultured on Matrigel-coated 6-well plates. Media was changed every other day, and cells were split using ReLeSR about once per week.

### Generation of iPSC-derived iMGLs

We followed the protocol as described in Abud et al. 2017^31^. iPSCs were first differentiated to hematopoietic stem cells over a period of 12 days. Briefly, iPSCs were dissociated into a single-cell suspension using TrypLE Select and plated at ∼500k cells/well onto tissue culture-treated 6-well plates in StemFlex + rock inhibitor. After 24 hours (HPC diff: day 1), the media and floating cells were collected, spun, and resuspended in differentiation medium (DM) 1, which consisted of HPC base medium supplemented with FGF2, BMP4, Activin A, LiCl, and rock inhibitor, and replated and cultured under hypoxic (5% O2) conditions. On day 3, cells were switched to DM 2, which included FGF2 and VEGF. On day 5, cells were switched to DM 3, which included FGF2, VEGF, IL6, TPO, SCF, and IL3, and switched back to a normoxic incubator. Cells were supplemented with additional DM3 every other day until floating HPCs were collected between days 11-15 and either frozen in Bambanker Freezing Medium or cultured directly in iMGL medium as described below.

HPCs were plated at 166k/well in a 6-well plate in iMGL medium, consisting of DMEM/F12, GlutaMAX, Non-essential amino acids, B27, N2, ITS, insulin, and MTG, and supplemented with fresh IL-34, M-CSF, and TGFb on each feeding day. Cells were fed every other day with 1mL medium, and were split 1:2 once per week over the course of the 25 day HPC → iMGL differentiation protocol. At day 25-28, cells were supplemented with the above factors along with two maturation factors: CX3CL1 and CD200. Cells were cultured in this medium for 3 days and used for subsequent assays on day 28-31. For RNA-seq and qPCR analysis: matured iMGLs were plated at 500k cells per well of a 6-well plate and treated with 10ng/mL of LPS or 20ng/mL of IFNγ. After 24 hours of treatment, cells were washed once with PBS and then lysed directly in the well using RLT buffer with beta-mercaptoethanol (Qiagen RNA-easy Mini Kit) and lysed cells were kept at −80C and processed together. Each staggered HPC → iMGL differentiation is considered one biological replicate.

### Patient recruitment

This study is part of the MyND (Myeloid cells in Neurodegenerative Diseases) initiative which aims to characterize the immune system (monocytes and microglia) from PD, Alzheimer’s disease and age-matched controls. For this study, we have selected a subset of samples including Controls and PD, with and without the G201S variant in the *LRRK2* gene. Sample collection was performed as part of ongoing research studies at two centers: Bonnie and Tom Strauss Movement Disorder Center at Mount Sinai Beth Israel (MSBI) and Fresco Institute for Parkinson’s and Movement Disorders at New York University (NYUMD). All participants gave informed consent at the respective institutions to contribute samples to this research, and participants included diagnosed PD cases, family members who harbored *LRRK2* G2019S variants, but did not have PD, and non-variant controls.

#### Mount Sinai/Mount Sinai Beth IsraelMovement Disorder (BPMD)

Diagnosis of PD was determined by movement disorder neurologist, when participants met the UK Parkinson’s disease society Brain Bank (UKBB) criteria for probable PD^61^, except that family history might be present.Family members participated in genetics research studies who both harbored and did not have the LRRK2 G2019S variant, and did not have known neurodegenerative diseases or auto-immune diseases were collected as controls. Participants were all co-enrolled in Genetics of Parkinson Disease Study and most in U01NS107016-01A1.

#### New York Movement Disorder (NYUMD)

Inclusion criteria followed the United Kingdom Parkinson’s Disease Society Brain Bank Clinical Diagnostic Criteria^62^. A population of age and sex-matched non-affected subjects was enrolled as controls, which did not have known PD at the time of evaluation and no history of any other relevant neurodegenerative disorder.

### PBMC isolation

All experiments were performed with monocytes derived from cryopreserved PBMCs to avoid any technical variability. Blood processing was performed as described before^7^. Briefly, blood was collected in Vacutainer blood collection tubes with acid citrate dextrose (ACD) (BD Biosciences) and was shipped to the Raj laboratory to be processed within the next 2-3 hours. Blood was centrifuged at 1,500 g for 15 mins, and aliquots of whole blood and plasma were stored at −80 °C for subsequent analysis. Secondly, SepMate tubes were filled with 15 ml Ficoll-Plaque PLUS (GE Healthcare) and 2-fold PBS (Gibco) diluted blood was added and centrifuged 15 mins at 1,200 g. After centrifugation, cells were washed with PBS and cryopreserved in 90% FBS (Germini) + 10% DMSO (Sigma Aldrich) at a concentration of 10 million cells/ml in Nalgene cryogenic vials (ThermoScientific). Vials were placed in NalGene CryoFreezing containers at −80 °C during 24-72 hours, and subsequently placed at liquid nitrogen for storage long-term.

### Monocyte isolation, culture and stimulation

Experiments were performed in 2 independent experimental batches. Monocytes were isolated from cryopreserved PBMCs as follows. Cryopreserved PBMCs were thawed in a water-bath and diluted in 10 ml of media (RPMI + 10% FBS + P/S + Gln) and centrifuged. After washing, PBMCs were sorted to monocytes using CD14^+^ magnetic beads (Miltenyi) in the Manual sorter following the manufacturer’s instructions. Monocyte number and viability was assessed using Countess II Automated Cell counter (Thermo Fisher). After sorting, monocytes were resuspended in supplemented RPMI media at a concentration of 1 million cells/ml. 500,000 cells were plated with the following stimuli: LPS-EK ultrapure (InvivoGen) 10 ng/ml, IFNγ (R&D Systems) 20 ng/ml or baseline (media + PBS). After 24 hrs of culture at 37 °C and 5% CO2, cells were washed with PBS and pellets were resuspended in 350 µl of RLT + 1% 2-Mercaptoethanol (Sigma Aldrich) and stored at −80 °C.

### RNA isolation and RNAseq

RNA was isolated using RNeasy Mini kit (Qiagen) following the manufacturer’s instructions and including the optional DNase treatment. RNA was stored at −80 °C prior to library preparation. Library preparation and sequencing was performed at Genewiz Inc. using the SMART-Seq v4 Ultra Low input library preparation protocol, which uses poly-A selection. Samples were sequenced with a depth of 30 million 150-bp paired-end reads using Illumina HiSeq 4000 platform.

### RNAseq analysis

#### RNA-seq data processing, quality controls and normalization

RNAseq data was processed as described before^7^. In brief, RAPID-nf was used to process FASTQ files^63^. After adapter trimming with trimmomatic (v0.36)^64^, STAR (2.7.2a)^65^ was used to align samples to the hg38 build (GRCh38.primary_assembly) of the human reference genome with indexes created from GENCODE (v30). For gene expression quantification, we used RSEM (1.3.1)^66^. Sequencing quality and technical metrics were assessed with FASTQC before alignment, and Picard (2.20) and Samtools (v1.9)^67^ post-alignment. No samples were removed based on these metrics. Multidimensional reduction using principal component analysis (PCA) and multidimensional scaling (MDS) before and after regressing covariates were used for the identification of outliers. No outliers were removed based on multidimensional reduction. Samples mismatches were identified by the expression of the sex genes *UTY* and *XIST* compared to the reported sex. One sample was removed because of sex mismatch.

Individual gene counts and transcripts per million (TPM) used for downstream analyses were generated using RSEM and assembled to a matrix via the tximport R package. Then, counts per million (CPM) were calculated using cpm() function from the edgeR packing in R. Lowly expressed genes were filtered out, which were defined as having less than one count per million in at least 10% of the samples.

#### Covariate selection using variancePartition

In order to avoid confounders, we used the R package variancePartition^68^to understand the major sources of variation in gene expression. Gene counts were normalized using TMM (trimmed means of M-values) calculated from edgeR, followed by voom transformation, which was used as input for variancePartition. Based on the variancePartition results we selected the covariates which explain the most variance in gene expression to be regressed out in the downstream analysis. For the iMGLs the covariates selected were: *% reads aligned, % of duplication + batch of RNA isolation*

For the human monocytes dataset we controlled variation as follows: *% coding bases + median 3-prime bias + gender + RNA concentration + age + % intergenic bases + the first 4 MDS from the genotype data*.

#### Differential gene expression analysis

For doing differential expression we used the R package limma^69^. The inputs included the count matrix and the covariate file. The data was normalized using TMM values calculated from edgeR, followed by voom transformation. P-values were adjusted for multiple comparisons using the Benjamini-Hochberg false discovery rate (FDR) correction.

#### Pathway enrichment analysis

Pathway analysis was performed using GSEA^70^ focusing on biological processes from GO. The input gene sets were generated ranking genes obtained by differential expression in terms of t-value. We used either the GSEA software or the fgsea function in R. We focused on biological processes from GO. To test specific pathways we used several curated gene lists from the bibliography (**Supplementary Table 11**). We tested the statistical enrichment using Fisher’s exact test.

In order to explore if genes were enriched for specific transcription factors, we used DoRothEA ^71,72^. We filtered regulons filtered at confidence levels “A” and “B” and performed enrichment using fGSEA with preranked data in terms of t value. Only regulons with adjusted p-value < 0.05 were considered.

### DNA isolation and genotyping

DNA was isolated using 1 ml from whole blood using the QiAamp DNA blood Midi kit (Qiagen) following the manufacturer’s instructions. DNA concentration and quality was assessed using nanodrop. For genotyping we used the Illumina Infinium Global Screening Array (GSA). Genetic and Ashkenazi Jewish ancestry was assessed as described before^7^. Additionally, targeted genotyping for specific regions was performed. G2019S variant at the *LRRK2* was outsourced and genotyped in Dr. William Nichol’s laboratory at the Cincinnati Children’s Hospital. Additionally, we also genotyped for the 11 most common GBA variants and for the 3 major APOE isoforms^7^.

### Immunocytochemistry and cell imaging

Cells were grown on 1.5mm coverslips coated with matrigel (iMGLs). Cells were first fixed in a 4% formaldehyde (Electron Microscopy cat. # 15714) solution in 4% sucrose in PBS (Thermo Fisher Scientific) for 10 minutes at room temperature, then washed twice with PBS and fixed with ice cold methanol (Sigma) for 10 minutes on ice. Cells were again washed twice with PBS and blocked using 2% normal donkey serum (NDS) (Thermo Fisher Scientific) with 0.1% triton-X (Sigma) for permeabilization for 1 hour at room temperature. Primary antibodies were diluted in 0.2% NDS and left on overnight at 4 C. AlexaFluor secondary antibodies were used at 1:500, and cells were mounted on coverslips using Vectashield hard-set mounting medium with DAPI. The following primary antibodies were used: anti-mouse IBA1 (FUJIFILM Wako, #019-19741), anti-Rabbit LRRK2 (Abcam, #ab13474).

### Cytokine/chemokine proteome assay

iMGLs were treated with IFNγ or LPS for 24 hours and cell supernatant was collected and frozen at −80 °C. Undiluted samples were sent to Eve Technologies and 48 cytokines were measured using the Human Cytokine/Chemokine Panel 48-Plex Discovery Assay Array (HD48A).

### Human myelin isolation

Myelin debris were isolated from frozen human post-mortem brains using the protocol previously described^73^. A pool of brain pieces from 4 control donors were mixed together to avoid any effect dependent on the individual. Brain tissue was homogenized in 0.32 M sucrose with protease inhibitors (Cell Signaling Technology) and sonicated for 10 mins. A sucrose gradient was prepared using 15ml of 0.85M sucrose and 15ml of 0.32M sucrose containing tissue homogenate and was centrifuged for 40 min at 24,4000 rpm using a Beckman L70 ultracentrifuge, with low acceleration and deceleration. Afterwards, the layer containing myelin was collected, washed with MilliQ water and centrifuged at 24,4000 rpm for 15 min. The pellet was submitted to osmotic shock by resuspending in water and centrifuging at 9800 rpm for 15 min twice. The pellet was then resuspended in 0.32M sucrose and combined with 0.85M sucrose in a 1:1 ratio, and centrifuged at 24,4000 rpm for 40 min. The interface was again collected and resuspended in 0.1 M sodium bicarbonate (Sigma-Aldrich). The volume was adjusted to obtain a final protein concentration of 1 μg/ml. Absence of contamination by endotoxins were assessed using a commercial kit following manufacturer instructions (GenScript) and myelin fragments were conjugated with 1:100 pHrodo red dye (Invitrogen) according to the manufacturer’s protocol and stored at −80 °C.

### Phagocytosis assays

Phagocytic capacity of iMGLs was measured using either red fluorescent carboxylate-modified polystyrene latex beads (Millipore Sigma, 0.9 μm) or pHrodo labeled human myelin. iMGLs were plated at ∼30,000 cells/well in a matrigel-coated 96-well plate and allowed to attach for 24 hours. Cells were then treated with fresh media alone, or media containing 20ng/mL of IFNγ or 10ng/mL of LPS for 24 hours. Fresh media was changed the next morning and cells were then treated with carboxylated beads or pH-rodo labeled human myelin for 24 hours. Cytochalasin D 2uM was added as negative control for the final 3 hours.

For myelin phagocytosis, phagocytosis was measured by imaging using IncuCyte by monitoring the plates during the 24 hrs before and after myelin addition. Pictures were taken every hour. Phagocytosis was assessed quantifying red object intensity and normalizing against phase confluence using the IncuCyte ZOOM software as described in detail^72^.

For assessing carboxylated-beads uptake, cells were washed once with 1x DPBS at the conclusion of treatment, and incubated in Live/Dead Violet stain (Sigma-Aldrich) for 15 minutes at 37 C. Cells were then washed again using 1x DPBS and resuspended in 140uL DPBS for flow cytometry using the Attune. After selecting viable cells, the phagocytic index was calculated as the percentage of red fluorescent-positive cells multiplied by the geometric mean of red fluorescence intensity divided by 10^6^. All treatments were performed with at least 3 technical replicates (each technical replicate as one well of a 96-well plate) and 3 biological replicates (each biological replicate as one differentiation).

### Proliferation capacity

In order to assess the proliferation capacity, cells were monitored for 24 hrs using IncuCyte. Proliferation of cells was quantified as phase confluence using IncuCyte ZOOM software. Experiments were performed with at least 3 technical replicates (each technical replicate as one well of a 96-well plate) and 3 biological replicates (each biological replicate as one differentiation).

## Data availability

The raw RNA-seq data from induced microglia-like cells (iMGLs) can be accessed on GEO (GSE240907), while stimulated monocyte data is available on dbGAP (study accession ID: phs002400.v1.p1) through this link: https://www.ncbi.nlm.nih.gov/projects/gap/cgi-bin/study.cgi?study_id=phs002400.v1.p1.

## Acknowledgements

We thank the study participants for providing blood samples. We thank Dr. Mark Cookson for sharing homozygous and heterozygous G2019S iPSC lines. T.R is supported by grants from Michael J. Fox Foundation (Grant #14899 and #16743) and the National Institutes of Health (NINDS U54-NS123743, NINDS R01-NS116006, NINDS U01-NS107016, NIA R21-AG063130, NIA R01-AG054005, NIA U01-AG068880, NIA RF1-AG065926, NIA R56-AG055824, NIA P30-AG066514). E.N. is supported by the Spanish Ministry of Science and Innovation (PID2022-139936OA-I00), the department of Genetics and Genomic Sciences at Mount Sinai Hospital (10-GGSPP19) and Universidad Complutense de Madrid. A.G.E was supported by the National Institutes of Health (NIA F31-AG059337)

## Author Contributions

EN, TR, and AGE conceived the study and led the project. EN and AGE designed and performed experiments and analysis, and wrote the manuscript. MP, AM and AA performed experiments and generated data. RAV, JH, BM, KL and EU helped in data analysis. DR, RS and MR recruited participants and GR, RS-P, SBB, VK, VLS, JM, KL and SF recruited and examined participants and determined status, and CA was in charge of clinical coordination. EN, TR, AMG and RS-P provided funding. TR, EM and AMG helped with intellectual discussion and interpretation of results. All authors reviewed and edited the manuscript.

## Notes

### Competing Interest Statement

The authors have declared no competing interest.

