## Supplementary Figures for "*LRRK2* G2019S variant is associated with transcriptional changes in Parkinson’s disease human myeloid cells under proinflammatory environment"

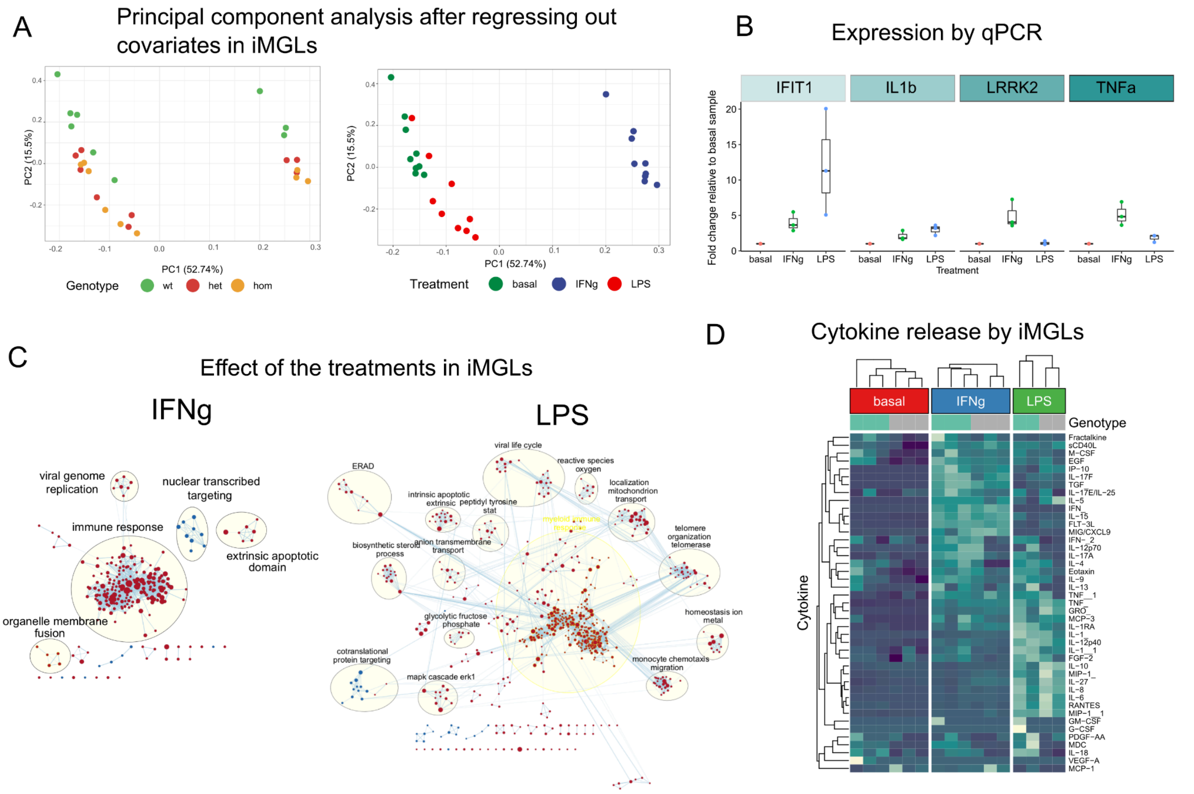


**Supplementary Figure 1. Human induced microglial cells (iMGLs) transcriptomic and cytokine analysis.** (A) Principal component analysis (PCA) of the 27 iMGL samples after regressing out technical covariates. Each dot represents a sample and is colored by genotype (left) or treatment (right). (B) qPCR validation of the expression of IFIT1, IL-1b, LRRK2 and TNFa in iMGLs (n=3). (C) Visualization of pathway dysregulation upon IFNγ (left) and LPS (right) treatment using Cytoscape. (D) Heatmap showing cytokine and chemokine levels assessed in iMGLs media. The heatmap shows the cytokines and chemokines (y axis) across the samples (x axis) clustered by treatment and genotype. Color represents the levels of expression.

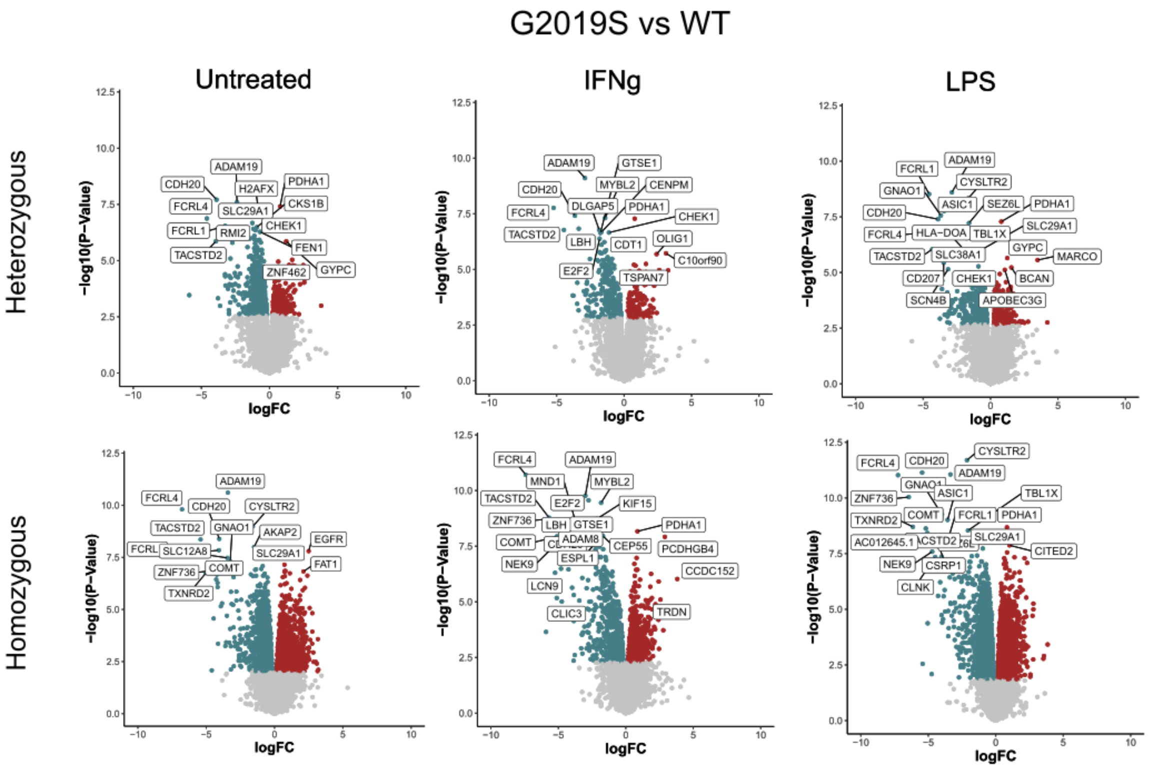


**Supplementary Figure 2. Transcriptomic dysregulation in G2019S het and hom vs WT iMGLs) transcriptomic.** Volcano plots showing differential expression between G2019S$^{+/-}$ (upper plots) or G2019S$^{+/+}$ (bottom plots) and WT under baseline (left), IFNγ (medium) or LPS (right). Plots show the FC of genes (x axis, $\log_2$ scale) and their *P* value significance (y axis, $-\log_{10}$ scale). DEGs at FDR < 0.05 are highlighted in red (upregulated genes) and blue (downregulated genes). A moderated *t*-statistic (two-sided) was used as the statistical test.

A

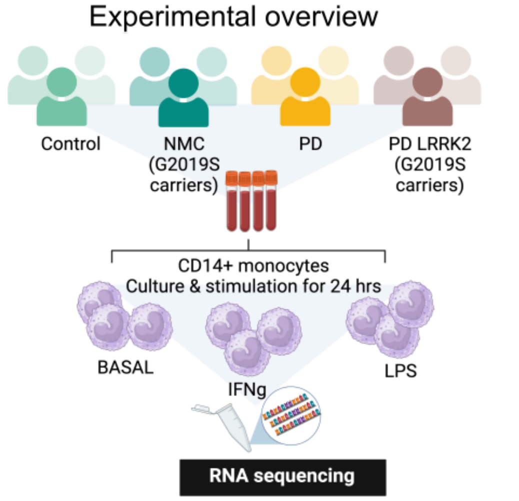


C

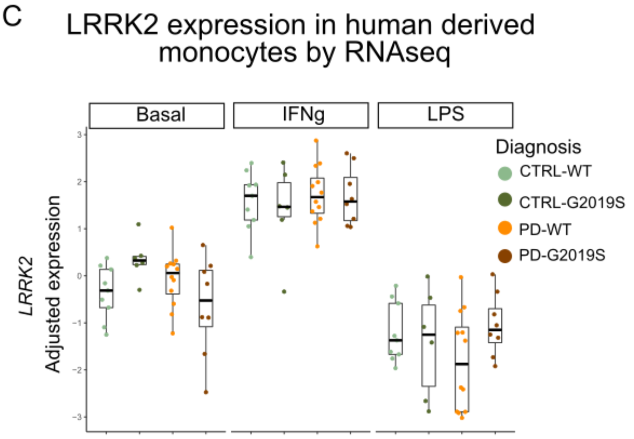


B

Demographic data

| | Donors | Sex | | Age |
|---|---|---|---|---|
| | #Number | F | M | Average |
| CTRL-WT | 10 | 7 | 3 | 65 |
| CTRL-G2019S | 6 | 3 | 3 | 67.2 |
| PD-WT | 12 | 4 | 8 | 64.6 |
| PD-G2019S | 8 | 4 | 4 | 68.2 |

D

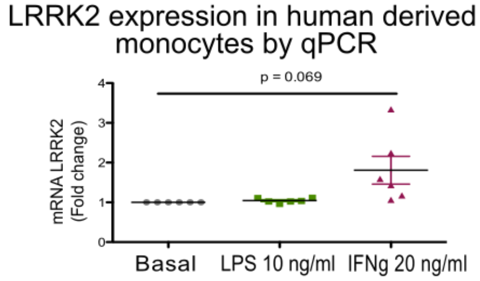


**Supplementary Figure 3. Human derived monocytes transcriptomic analysis.** (A) Experimental workflow for the generation of monocyte transcriptomic profiles. (B) Tables describing the samples included in the study. (C) Expression of LRRK2 in human derived monocytes upon the different treatments in Controls (n=10), NCM (n=6), PD (n=12) and PD-LRRK2 (n=8). Adjusted gene expression levels after normalization are shown. (D) qPCR validation of LRRK2 expression in control monocytes upon the different stimulations (n=6). Boxplots: the line represents the median. The boxes extend from the 25th - 75th percentile.

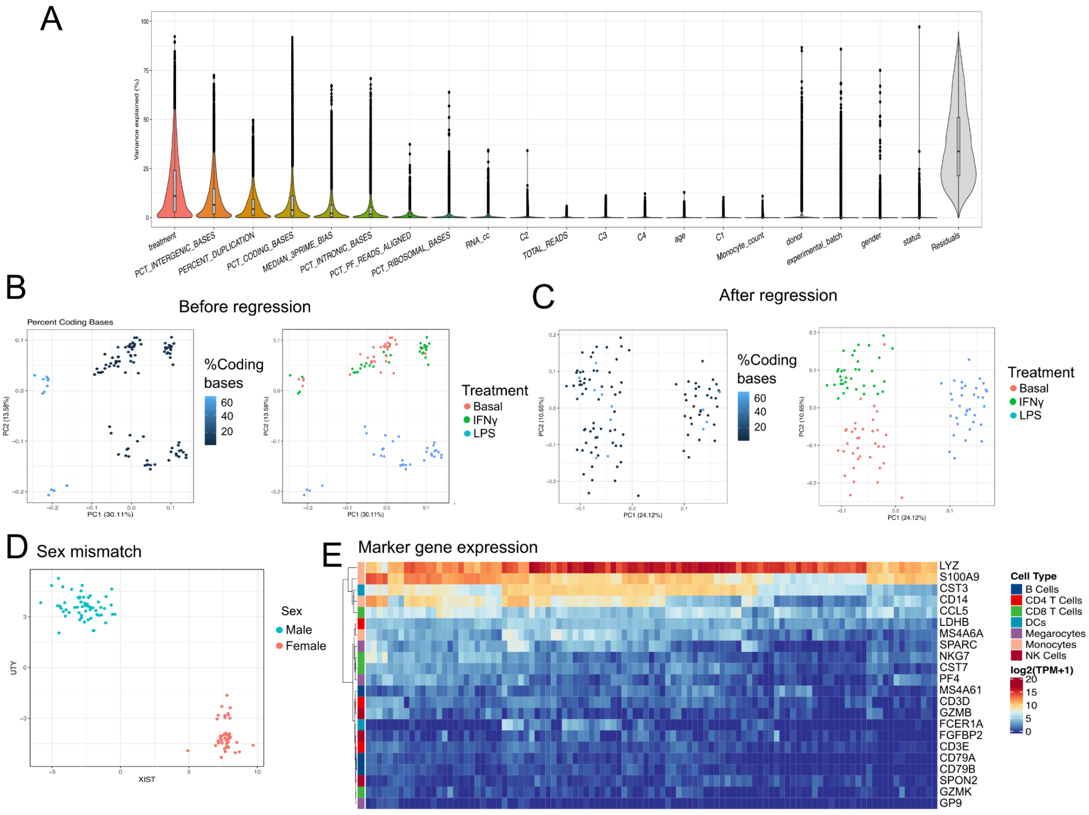


**Supplementary Figure 4. Quality control in monocyte transcriptomic analysis.** (A) Violin plot showing the % of variance (y axis) explained by known technical and biological covariates (x axis) by variancePartition. (B) Principal component analysis (PCA) of the samples (105 samples from 35 independent donors) before and (C) after regressing out covariates. Each dot represents a sample. Samples are colored by percent of coding bases (left) and treatment (right). (E) PCA showing the expression of the male marker *UTR* and the female marker *XIST*. Samples are colored by sex, showing no sex mismatches. (F) Heatmap of the expression ($\log_2$(TPM +1)) of marker genes of different blood cell types (represented by colors). Gene markers are represented in the y axis and samples in the x axis (n=105).

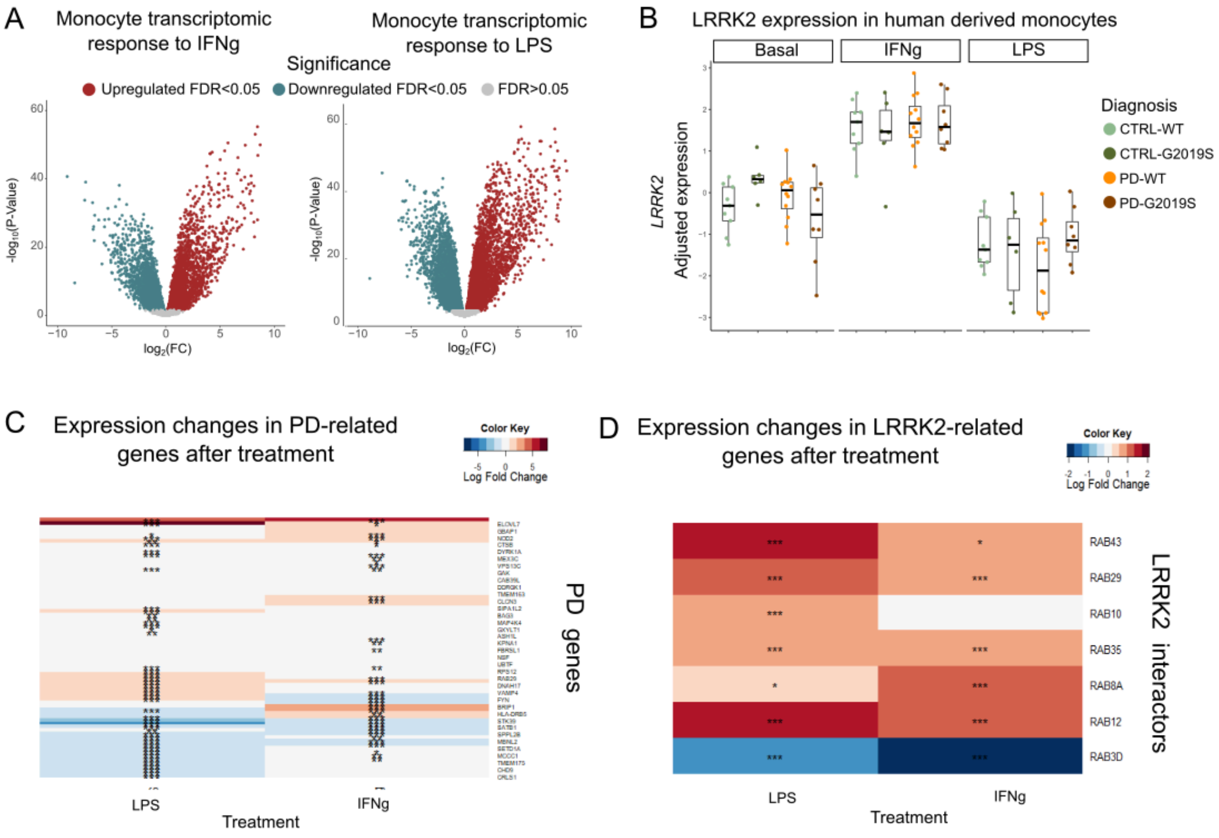


**Supplementary Figure 5. Transcriptomic dysregulation in human monocytes upon IFNγ and LPS stimulation**. (A) Volcano plot showing differential expression between IFNγ and (B) LPS stimulation. Plots show the FC of genes (x axis, $\log_2$ scale) and their *P* value significance (y axis, $-\log_{10}$ scale). DEGs at FDR < 0.05 are highlighted in red (upregulated genes) and blue (downregulated genes). A moderated *t*-statistic (two-sided) was used as the statistical test. (C) Heatmap showing the logFC of genes associated with PD or (D) LRRK2 interactors. The heatmap shows a selection of genes (y axis) across samples (x axis), clustered by genotype. Blue represents logFC < 0 and red represents logFC > 0.

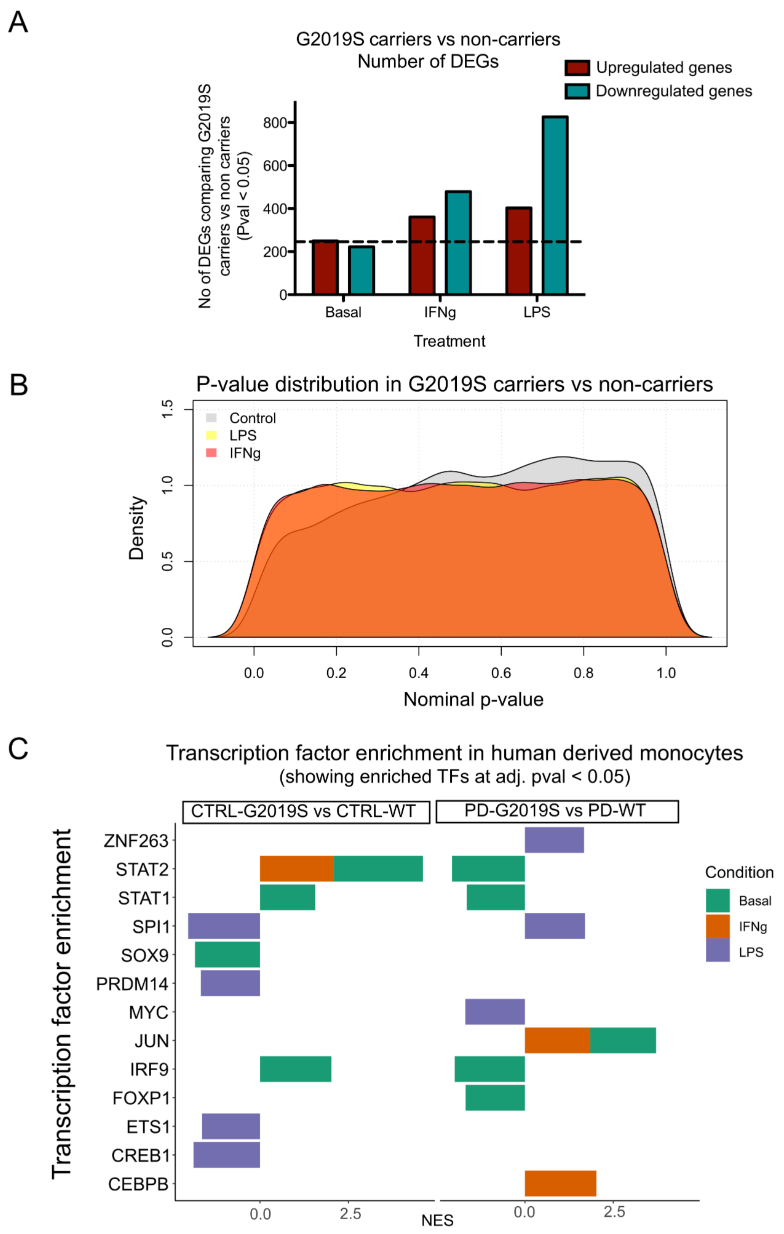


**Supplementary Figure 6. Transcriptomic dysregulation in G2019S carriers upon diagnosis in IFNγ and LPS stimulation**. (A) Boxplot showing the number of DEGs at nominal p-value between G2019S carriers and non carriers. Red plots represent the number of up-regulated genes and blue down-regulated genes. (B) Density plot showing the p-value distribution of G2019S carriers vs non-carriers upon the different stimulations (represented by colors). (C) Transcription factor enrichment in human derived monocytes of CTRL-G2019S vs CTRL-WT (left) and PD-G2019S vs PD-WT (right) upon the different stimulations (represented by colors). Transcription factors are expressed in the y axis and normalized expression score in the x axis. Only regulons significantly enriched at a q-value < 0.05 are shown.

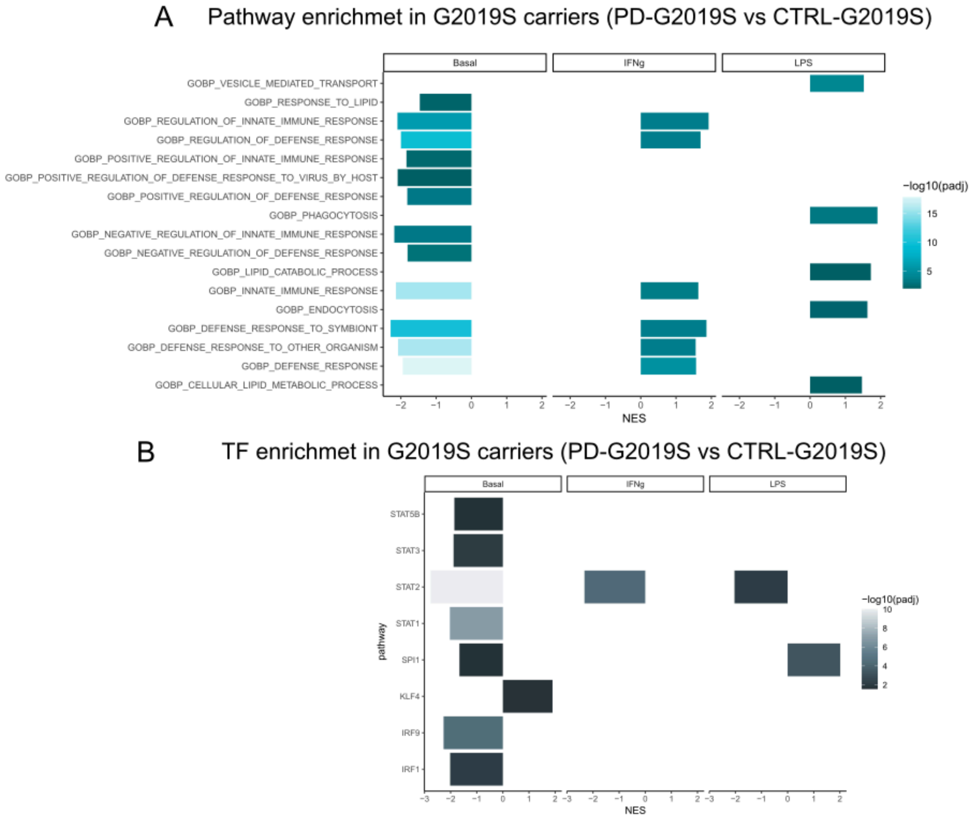


**Supplementary Figure 7. Pathway and TF enrichment analysis in PD-G2019S vs CTRL-G2019S in untreated and upon IFNγ and LPS stimulation**. (A) Pathway enrichment analysis of PD-G2019S vs CTRL-G2019S. Pathway enrichment was performed with a pre ranked approach from GSEA using Biological Processes. Y axis shows the significant pathways and x axis normalized expression score. Only a subset of significant pathways are represented (full data is included in Supplementary Table 35. (B) Transcription factor enrichment in human derived monocytes of PD-G2019S vs CTRL-G2019S upon the different stimulations. Transcription factors are expressed in the y axis and normalized expression score in the x axis. Only regulons significantly enriched at a q
